# *RRAS* and *RRAS2* mutations are recurrent oncogenic drivers in lung cancer and are sensitive to the pan-RAS inhibitor RMC-6236

**DOI:** 10.1101/2025.09.19.677259

**Authors:** Alexander J. Pfeil, Tom Zhang, Ryan Cheng, Marissa S. Mattar, Juan Luis Gomez Marti, Leo Gili, Rachel Lai, Inna Khodos, Rohan P. Master, Jenna-Marie Dix, Andrea Gazzo, Kathryn C. Arbour, Elisa de Stanchina, Christopher A. Febres-Aldana, Marc Ladanyi, Soo-Ryum Yang, Romel Somwar

**Affiliations:** Department of Pathology and Laboratory Medicine, Memorial Sloan Kettering Cancer Center, New York, NY, 10065; University of North Carolina School of Medicine, Chapel Hill, NC, 27599; Graduate Medical Education, Memorial Sloan Kettering Cancer Center, New York, NY, 10065; New York Medical College, Valhalla, NY 10595; Human Oncology and Pathogenesis Program, Memorial Sloan Kettering Cancer Center, New York, NY, 10065; Department of Pathology and Laboratory Medicine, Northwell Health Lenox Hill Hospital, New York, NY, 10075; Anti-tumor Core Facility, Memorial Sloan Kettering Cancer Center, New York, NY, 10065; Thoracic Oncology Service, Department of Medicine, Memorial Sloan Kettering Cancer Center, New York, NY, 10065; Laboratory of Pathology, National Cancer Institute, National Institute of Health, Bethesda, MD, 20892, USA

**Author notes:** These authors contributed equally to this manuscript (Alexander J. Pfeil, Tom Zhang). Co-senior authors (Soo-Ryum Yang, Romel Somwar). **Author for Correspondence: Marc Ladanyi, MD**, Department of Pathology and Laboratory Medicine, Memorial Sloan Kettering Cancer Center, 1275 York Ave, NY, NY, 10065.

## Abstract

**Introduction:** *RRAS* and *RRAS2* encode a subfamily of RAS-like small GTPases that share considerable structural and functional similarities with *KRAS*, *HRAS*, and *NRAS.* Whether homologous *RRAS*/*RRAS2* mutations are oncogenic and actionable drivers in lung cancer remains underexplored.

**Methods:** An institutional cohort of 8,488 non-small cell lung carcinomas (NSCLC) sequenced by comprehensive targeted DNA sequencing (MSK-IMPACT) between 2016-2024 was evaluated for *RRAS*/*RRAS2* mutations. *RRAS*^Q87L^ or *RRAS2*^Q72L^ were modeled in murine IL3-dependent Ba/F3 cells and immortalized human bronchiolar epithelial cells (HBECs). The oncogenic potential, signaling characteristics, and sensitivity to PI3K and MAPK pathway inhibitors, including the novel pan-RAS inhibitor RMC-6236, were evaluated *in vitro* and *in vivo*.

**Results:** *RRAS*^Q87L^ or *RRAS2*^Q72L^, homologous to KRAS-codon Q61 substitutions, were found in ∼0.45% of NSCLCs (38/8,488), with all but two lacking other MAPK pathway oncogenic drivers. RRAS^Q87L^ and RRAS2^Q72L^ mutations transformed Ba/F3 and HBEC cells and robustly activated MAPK and PI3K-mTOR pathway signaling. RMC-6236 suppressed proliferation of RRAS^Q87L^ and RRAS2^Q72L^ mutant cell lines, reduced ERK phosphorylation, induced apoptosis, and impeded cell-cycle progression. *In vivo*, RMC-6236 significantly inhibited growth of *RRAS*^Q87L^/*RRAS2*^Q72L^-mutant HBEC-derived xenografts.

**Conclusions:** RRAS^Q87L^ and RRAS2^Q72L^ are recurrent, oncogenic, and potentially actionable drivers in NSCLC. Our study supports the inclusion of *RRAS/RRAS2* into routine molecular diagnostic panels for precision oncology and provides preclinical rationale for investigating the potential therapeutic utility of pan-RAS inhibitors for patients with RRAS^Q87L^/RRAS2^Q72L^-mutant lung cancers.

**Statement of translational relevance:** Targeted therapies have transformed standard of care for oncogene-driven non-small cell lung carcinomas (NSCLC), yet a significant subset lacks actionable drivers. We identified recurrent *RRAS*^Q87L^ and *RRAS2*^Q72L^ mutations which are mutually exclusive with other MAPK pathway drivers and found in ∼0.45% of NSCLC, comparable in prevalence to *NTRK* and *NRG1* fusions. In preclinical models, these mutations activate canonical growth signaling, drive tumorigenic phenotypes, and confer sensitivity to RAS/MAPK-directed agents, including the novel pan-RAS inhibitor RMC-6236, currently in trials for patients with solid tumors harboring *KRAS* mutations. These data support *RRAS*^Q87L^ and *RRAS2*^Q72L^ as *bona fide* lung cancer drivers and nominate RRAS/RRAS2-mutant tumors as candidates for pan-RAS-targeted therapeutics. Our findings provide a biologic rationale and preclinical evidence to inform molecular testing paradigms and to prioritize enrollment of patients with RRAS/RRAS2-mutant NSCLC into future clinical trials of pan-RAS inhibitors.

## INTRODUCTION

The *RAS* family of small-GTPases are among the most frequently mutated oncogenes in human cancers, present in approximately 10-30% of all tumors.^1,2^ These “molecular switches” alternate between their GDP-bound, inactive state and GTP-bound, active state in response to growth factor-receptor-tyrosine kinase (RTK) signaling to regulate cellular growth, proliferation, and survival.^3^ Hyperactivation of RAS family members due to genetic mutations is capable of driving tumorigenesis via the RAF/MEK/ERK (MAPK) pathway.^4^ The most well-studied RAS isoforms are KRAS, HRAS, and NRAS, all sharing roughly 80% primary amino acid sequence homology.^5^ *KRAS* is the most frequently mutated oncogene in most tumor types and is particularly prevalent in colon, lung and pancreatic cancers.^6^ In lung adenocarcinoma (LUAD), *KRAS* mutations are seen in up to a third of tumors, with single amino acid substitutions at codons 12, 13 and 61 being the most common hotspots.^7^ Small molecule inhibitors of KRAS^G12C^, including sotorasib and adagrasib, have demonstrated clinical efficacy and proof-of-concept for targeting mutant KRAS in multiple cancer types.^8^ These therapies, however, are currently restricted to KRAS^G12C^-expressing tumors, which represent only 40% of KRAS mutant non-small cell lung carcinomas (NSCLCs) and 10-13% of all NSCLCs.^9^ The other RAS isoforms are also important drivers of tumorigenesis found in 25% of all RAS-mutant cancers.^2^ As such, investigation into targeting additional RAS mutants and inhibitors that target all RAS family members with broader applicability has accelerated, with multiple promising therapies now under clinical investigation.^10^

*RRAS* and *RRAS2* encode members of a distinct but related subfamily of RAS-like GTPases that share approximately 55% sequence identity with classical RAS isoforms. Although highly conserved in regions involved in guanine nucleotide-binding, GEF/GAP interactions, and effector binding, RRAS and RRAS2 possess unique N- and C-terminal regions that influence subcellular localization and differential activation of downstream effectors.^11^ Hence, RRAS proteins are thought to have both overlapping and distinct signaling properties compared to classic RAS-family proteins, capable of activating RAF, PI3K, and RAL-GEF amongst other effectors, to promote cellular proliferation and survival.^11^ RRAS primarily localizes to focal adhesions and is involved in inside-out activation of integrin signaling, contributing to the balance of migration and adhesion signaling.^12^ In addition to activating the MAPK pathway, some reports have suggested that RRAS can preferentially signal through the PI3K-AKT pathway in certain cellular contexts.^13,14^ For instance, constitutively activated RRAS in breast epithelial cells induced migration through PI3K activation, but not MAPK.^15^ RRAS2 is also capable of activating both MAPK and PI3K-AKT signaling, albeit with weaker affinity for RAF-family proteins and subsequently weaker ERK1/2 stimulation than other RAS-family members.^16,17,18^ In multiple cell types, RRAS2 has been demonstrated to signal in a MAPK-independent manner, with knockout models of B and T cells resulting in impaired activation of the PI3K-AKT-mTOR pathway. Hence, the primary signaling axis influenced by RRAS and RRAS2 appears to be cell-type and context specific.

Mutations in the regions of RRAS and RRAS2 involved in guanine nucleotide exchange or intrinsic GTPase activity have been shown to induce transformation *in vitro*.^19,20^ Many of these mutations occur at residues homologous to those at KRAS hotspots, primarily at codons 12 and 61, corresponding to codons 38 and 87 in RRAS and 23 and 72 in RRAS2, respectively. Furthermore, ectopic expression of wild-type RRAS2 is capable of transformation in pre-clinical models of triple-negative breast cancer and chronic lymphocytic leukemia.^21,22^ Despite homology with common driver genes and oncogenicity *in vitro,* few studies have identified *RRAS* or *RRAS2* mutations in human cancers. *RRAS2* mutations have been identified at high frequencies in *KIT*-mutated germ cell tumors (11.8%) and central nervous system germinomas (14.6%).^23,24^ Germline mutations in the *RRAS* family have also been identified in cases of “RASopathies”, a family of congenital disorders driven by the dysregulation of RAS signaling, including Noonan syndrome and associated juvenile myeloid neoplasms.^25^

Due to the rarity of *RRAS* and *RRAS2* mutations in clinical practice, the relevance and therapeutic implications of these mutations have not been well-studied. Moreover, the prevalence of *RRAS* family mutations in lung cancers remains poorly characterized, potentially owing to the lack of coverage of *RRAS* and *RRAS2* in most commercial and academic tumor DNA sequencing panel-based assays. By leveraging the MSK-IMPACT panel with complete coverage of *RRAS* and *RRAS2*, we were able to identify hotspot mutations within these two genes in NSCLCs that lacked other known activating alterations in the RTK-RAS-RAF pathway. In this study, we set out to characterize the signaling alterations caused by these mutations and more importantly, if these signaling programs render such tumors susceptible to targeted therapies including the novel and clinically active pan-RAS inhibitors given the sequence homology between RRAS/RRAS2 and the other RAS family members.

## MATERIAL AND METHODS

### Case selection

The study was approved by the Institutional Review Board (IRB) of MSKCC and conducted in accordance with the principles outlined in the Declaration of Helsinki. All patients in this study signed a written informed consent form a part of an IRB-approved protocol. A retrospective search for *RRAS* and *RRAS2* mutations was performed for NSCLC tissue samples that were genotyped using select MSK-IMPACT panels with full coverage of all *RRAS* (NM_006270) and *RRAS2* (NM_012250) exons, namely the MSK-IMPACT 468 and 505 panels between September 2016 to March 2024. In patients with multiple NSCLCs submitted for sequencing, the genomic profile from the initial tumor or those with *RRAS/RRAS2* mutations were included for analysis.

### MSK-IMPACT testing

Genomic DNA from tissue (tumor) and matched blood samples (normal) were extracted for MSK-IMPACT as described previously.^26^ In brief, MSK-IMPACT is a hybrid capture-based next-generation sequencing (NGS) assay that interrogates all exons and select introns of cancer-related genes (468 or 505-genes) to identify mutations, copy number alterations, and select structural variants from tumor tissue. The gene lists for the 468 and 505 panels are provided in **Supplementary Table 1**. Tumor mutation burden (TMB) was calculated by dividing the total number of nonsynonymous somatic mutations (single nucleotide and small insertions/deletions) by the size of the targeted coding regions. All somatic alterations that were (i) annotated as oncogenic and likely oncogenic by OncoKB^27^ and/or (ii) present in the cancer hotspot database^28^ were classified as driver/pathogenic; all else were considered variants of unknown significance. For all variants outside of *RRAS* and *RRAS2*, only driver alterations were included for analysis.

### Clonality estimation

Two methods were used to infer clonality of *RRAS* and *RRAS2* mutations. First, cancer cell fraction (CCF) for *RRAS*/*RRAS2* mutations was calculated in samples with available and evaluable FACETS data.^29^ Mutations with CCF >80% were classified as clonal. Secondly, in cases without FACETS data, relative VAF fraction (RVF) was used to estimate clonality and was calculated by dividing the VAF of *RRAS*/*RRAS2* mutations by the average of the VAFs from other non-*RRAS*/*RRAS2* mutations in a given sample. Mutations with RVF >80% were classified as clonal.

### Clinicopathologic data

Clinical parameters were extracted from the electronic medical records. All tumors underwent central pathologic review for histologic diagnosis according to the 2021 World Health Organization (WHO) classification.^30^ For invasive resected lung adenocarcinomas, comprehensive histologic subtyping was performed; using these histologic patterns, IASLC grades were determined.^31^

### PD-L1 immunohistochemistry

PD-L1 IHC (E1L3N; dilution 1:100, Cell Signaling Technologies, Danvers, MA) was performed on formalin-fixed paraffin-embedded tissue sections using an automated staining platform (Bond III, Leica). PD-L1 expression was measured using Tumor Proportion Score (TPS) as defined by the percentage of viable tumor cells with partial or complete membranous staining. PD-L1 positivity was defined as having a TPS score of ≥1%.

### Reagents and Cell Culture

All cell lines were cultured in a humidified incubator maintained at 5% carbon dioxide at 37°C. The human bronchiolar epithelial cell (HBEC) line was maintained in keratinocyte serum-free media supplemented with 5 ng/mL recombinant EGF and 50 µg/mL bovine pituitary extract (ThermoFisher, Waltham, MA). Untransformed Ba/F3 cells were maintained in RPMI-1640 supplemented with 10% FBS and 5 ng/mL murine IL3 from R and D Systems (Minneapolis, MN). Prior to Western blotting experiments, both cell lines were incubated with growth-factor free media for 24 hours prior to sample extraction or treatment. The presence of the mutations of interest within cDNA constructs was confirmed by polymerase chain reaction each time a cell line vial was thawed. Cells were tested for mycoplasma every three months and no cell line was tested positive throughout this study. RMC-6236 was purchased from STARK Chemicals (New York, NY) and prepared with DMSO for *in vitro* experiments. Primary antibodies used in Western blotting were purchased from Cell Signaling Technologies (Danvers, MA) or Proteintech (Rosemont, IL). Secondary anti-mouse or anti-rabbit IgG antibodies conjugated to horseradish peroxide were purchased from R and D Systems (Minneapolis, MN). See **Supplementary Table 2** for details of antibodies used in this study.

### cDNA Overexpression of Wild-type and Mutant RRAS and RRAS2

Nucleotides encoding RRAS WT, RRAS Q87L, and RRAS2 WT were ordered as gblocks from Integrated DNA Technologies (Coralville, IA). See **Supplementary Table 3** for exact sequences of gblocks. *RRAS* WT and *RRAS* Q87L sequences were partially codon optimized with no change in predicted amino acid sequence (NM_006270 and its mutated Q87L version), respectively. RRAS2 WT is identical to the coding sequence from the refseq NM_012250. The gblocks and the pCX4 bleo empty vector were digested with AgeI and EcoRI FastDigest enzymes (ThermoFisher, Waltman, MA) for 1 h at 37°C. Bands corresponding to the expected size of the digested products were resolved on 1% (wt/vol) agarose gel following gel electrophoresis at 130 V for 1 h and then extracted using QIAGEN’s QIAquick Gel Extraction Kit (Germantown, MD). Assembly of the final plasmids was done utilizing a T4 ligase reaction (ThermoFisher, Waltman, MA) with a 3:1 ratio of insert:backbone. Stable competent E. coli (New England Biolabs, Ipswich, MA) were transformed, and plasmids were extracted using QIAGEN’s HiSpeed Plasmid Midi Kit. RRAS2 Q72L was generated with Agilent’s (Santa Clara, CA) QuikChange Lightning Site Directed Mutagenesis Kit along with RRAS2 forward (5’-tggctccaaactcttctagtcctgctgtatccaaa-3’) and reverse (5’-tttggatacagcaggactagaagagtttggagcca-3’) primers. The following PCR thermocycler conditions were utilized: initial denaturation (95°C, 2 min), followed by 18 cycles of denaturation (95°C, 20s), annealing (60°C, 10s) and extension (68°C, 3 min 30s); the reaction was finished after the final extension step (68°C, 5 min). XL10 Gold Ultracompetent cells (Agilent, Santa Clara, CA) were transformed, and plasmids extracted using QIAGEN’s HiSpeed Midiprep Kit (Germantown, MD).

To generate virus with the cDNA of RRAS WT, RRAS Q87L, RRAS2 WT, and RRAS2 Q72L, HEK-293T cells were plated in 6 well plates at a density of 500,000 cells per well. The following day they were transfected with the corresponding cDNA along with pGp and pE-Ampho plasmids utilizing FuGENE® HD Transfection Reagent (Promega, Madison, WI) in a 3 uL FuGENE:1 ug plasmid ratio. Virus was harvested for three days, filtered with Millipore Steriflip® Vacuum Tube Top Filter, and then supplemented with polybrene 10 ug/mL (Millipore-Sigma, Burlington, MA). HBECs, and Ba/F3 cells were plated at a density of 250,000 and 1,000,000 cells, respectively, in 6 well plates prior to being infected with the above virus media supplemented with 100 umol HEPES and 5 ng/mL murine-IL3 (Ba/F3) or recombinant EGF (HBECs) and centrifuged at 37°C for 1 hr. These cells were treated with 750 µg/mL Zeocin^TM^ Selection Reagent (ThermoFisher, Waltham, MA) for 1 week following viral infection. For transient transfection of HEK-293T with the RRAS and RRAS2 plasmids, HEK-293Ts were plated at a density of 500,000 cells per well and transfected with just the RRAS and RRAS2 plasmids mixed with FuGENE® HD Transfection Reagent in a 3 uL FuGENE:1 ug plasmid ratio.

### Animal Studies

Mice were monitored daily and cared for following guidelines approved by the Memorial Sloan Kettering Institutional Animal Care and Use Committee and Research Animal Resource Center (protocol 04-03-009). Five million Ba/F3 or ten million HBEC cells mixed with 50% (vol/vol) matrigel were implanted via subcutaneous injection into the flank of 6-week-old female NOD-*scid* gamma (NSG) mice (RRID:IMSR_JAX:005557). To test if cell lines formed xenograft tumors cells were injected into two flanks of three mice per cell line (for a total of 6 tumors). For drug treatment studies, mice (five per group) were randomized by tumor size once tumor volume reached 100-150 mm^3^. Treatments included once daily (q.d.) doses via oral gavage of vehicle, 10 mg/kg RMC-6236, 25 mg/kg RMC-6236, or 50 mg/kg RMC-6236. RMC-6236 was reconstituted in 5% DMSO+40% PEG300+5% Tween80+50% ddH_2_O. Body weight and tumor size were determined two times per week, with tumor volume calculated with the formula: [length × (width)^2^]/2. Animals were monitored daily. Tumors were collected 3 hours after the last treatment.

### Cell Viability Assays

Cells were plated in 96-well clear bottom, white polystyrene plates at densities of 3,000 (Ba/F3) or 6,000 (HBEC) cells per well and treated with drugs for 96 hours. Quantification of viable cells was determined using alamarBlue viability dye (ThermoFisher, Waltham, MA) with fluorescence measured using a Molecular Devices SpectraMax M2 multimodal plate reader (Ex: 555 nm, Em: 585 nm). Data were analyzed with GraphPad Prism (RRID:SCR_002798) version 10 software by nonlinear regression. Curves were fitted to determine the concentration that inhibits 50% of cell viability (IC_50_). Data are expressed relative to control values with an average of two to four independent experiments in which each condition was assayed in triplicates.

### Western Blotting

For dose-response studies in HBECs, 500,000 cells were plated in 6-well dishes and 24 hours later cells were deprived of serum for 24 hours prior to treatment with inhibitors for 90 min. Cells were washed twice with ice-cold PBS and then lysed with 500 µL 2X LSB buffer supplemented with 7.5% beta mercaptoethanol, phosphatase and protease inhibitor (Selleckchem, Houston, TX), and 1 mM sodium orthovanadate (MilliporeSigma, Burlington, MA). Lysates were then passed through an insulin syringe to shear the DNA then heated to 55°C for 15 min. 10-20 µL of lysates were resolved by SDS-PAGE. For dose-response studies in Ba/F3 cells, two million cells were plated in 6-well plates and then deprived of serum for 3 hours. Cells were then treated with inhibitors for 90 min, collected by centrifugation and washed twice with ice-cold PBS. Cells were lysed in radioimmunoprecipitation assay (RIPA) buffer supplemented with phosphatase and protease inhibitors (Selleckchem, Houston, TX), and 1 mM sodium orthovanadate (MilliporeSigma, Burlington, MA). For time course studies, HBEC cells were serum-starved for 24 hours, then treated with 10 nM RMC-6236 for the indicated time periods. Inhibitor was added so that all treatments ended at the same time. Serum-free media with fresh inhibitor was added at each time point. Protein concentration was measured via Bradford assay (Bio-Rad, Hercules, CA). Twenty µg whole-cell lysates per sample were denatured with 1× Laemmli sample buffer for 10 minutes at 55°C and resolved by SDS-PAGE using 4%-12% Bolt gels (ThermoFisher, Waltham, MA). For *ex vivo* tumor sample westerns, available residual tumors post xenograft studies for vehicle and 25 mg/kg RMC-6236 treatment conditions were homogenized with an AgileGrinder^TM^ ACT-AG3080 (ACTGene, Piscataway, NJ) and then digested with RIPA buffer in the same manner as stated above for the Ba/F3 dose response studies. Ten µg of each *ex vivo* tumor sample was resolved by SDS-PAGE using 4-12% Bolt gels. For HEK-293T cells transiently transfected with the RRAS and RRAS2 plasmids, lysates were also prepared with RIPA buffer and ten µg lysate was resolved by SDS-PAGE using 4-12% Bolt gels. Separated proteins were next transferred onto polyvinylidene fluoride membranes and blocked in 3% (wt/vol) bovine serum albumin (Sigma, Marlborough, MA) solution for 1 hour at room temperature before addition of primary antibodies. Membranes were allowed to incubate overnight in primary antibody solution on a shaker at 4°C and then washed three times with Tris-buffered saline supplemented with 0.1% (vol/vol) Tween-20 (Santa Cruz Biotehnology, Santa Cruz, CA). Bound primary antibodies were detected with peroxidase-labeled goat antibody raised against rabbit or mouse IgG (R&D Systems, Minneapolis, MN). Enhanced chemiluminescence Western blotting detection reagent (Cytiva, Marlborough, MA) was added to the samples before capture on radiograph films. For just the *ex vivo* tumor sample western and HEK-293T cell line Western blot experiments, the blots were imaged with the ChemiDoc MP imaging system and Clarity ECL Western Blotting Substrates were utilized (Biorad, Hercules, CA). Western blot experiments were repeated at least two times with independently prepared protein samples.

### Western Blot Quantification

Densitometry values for the listed proteins were obtained using Fiji software. The phospho:total protein ratios were obtained by normalizing the phospho-protein and total protein densitometry values to their respective loading controls and then dividing them by each other. For the *ex vivo* tumor samples, graphs were made from the results of three independent tumors ran on the same gels.

### Statistical analysis

For *in vitro* IC_50_ determination, normalized values were plotted using a nonlinear regression fit to the normalized data with a variable slope. IC_50_ values were calculated by interpolating the concentration of the compound at which 50% inhibition occurred. 95% confidence intervals (CI) were used to compare efficacy. In animal models, area under curve analysis of tumor volume over time was used to compare treatment responses. For Westerns, values were compared via unpaired t tests when two groups were compared or with one-way ANOVA followed by Dunnett’s multiple-comparisons test for experiments with more than two groups. All tests of significance were two-tailed, and P < 0.05 was considered statistically significant. Error bars represent means ± standard error of mean (SEM).

## RESULTS

### RRAS and RRAS2 mutations in NSCLC

From September 2016 to March 2024, 8,488 patients with NSCLC were genomically profiled using MSK-IMPACT with full coverage of *RRAS* and *RRAS2* exons. In total, mutations in *RRAS* and *RRAS2* were detected in 40 and 41 patients, respectively. In *RRAS*, there was clear predominance of a hotspot missense mutation (c.260A>T, p.Q87L) that was present in 16 samples **(Figure 1A, top panel)**. All other mutations were observed only once. The RRAS Q87 residue is analogous to Q61 in classical RAS proteins, while RRAS G38 and G39 correspond to RAS G12 and G13, respectively. One case showed *RRAS* p.G38D, and another showed p.G39V. Alignment of the amino acid sequences of RAS isoforms are depicted in **Figure 1B**. In *RRAS2*, there was an equally notable hotspot mutation (c.215A>T, p.Q72L) observed in 22 samples **(Figure 1A, bottom panel)**. *RRAS2* p.Q72H was seen in one sample. As in *RRAS*, this hotspot mutation occurred in a residue analogous to Q61 in H/K/N RAS proteins **(Figure 1B)**. In contrast to Q72, analysis of G23 and G24 residues in RRAS2 (corresponding to G12 and G13 in classical RAS, respectively) showed only rare substitutions at low frequencies (p.G23C, n=1; p.G23V, n=1; p.G24C, n=2; p.G24V, n=1) similar to the background rates for variants of unknown significance. RRAS Q87 and RRAS2 Q72, as with KRAS Q61, are located in the switch II region common to RAS-family proteins, situated near the guanine nucleotide binding site **(Figure 1C and Supplementary Figure 1)**.^32^ KRAS mutations at the Q61 position block interactions with GTPase-activating proteins (GAPs) as well as intrinsic GTP hydrolysis, leading to hyperactivation.^33^ Taken together, the two dominant hotspot mutations, *RRAS* p.Q87L and *RRAS2* p.Q72L, were detected in 0.45% of patients with NSCLC (38/8,488). Given the strikingly recurrent nature of *RRAS*^Q87L^ and *RRAS2*^Q72L^ mutations, patient samples with these variants were further analyzed as separate cohorts.

**Figure 1.**
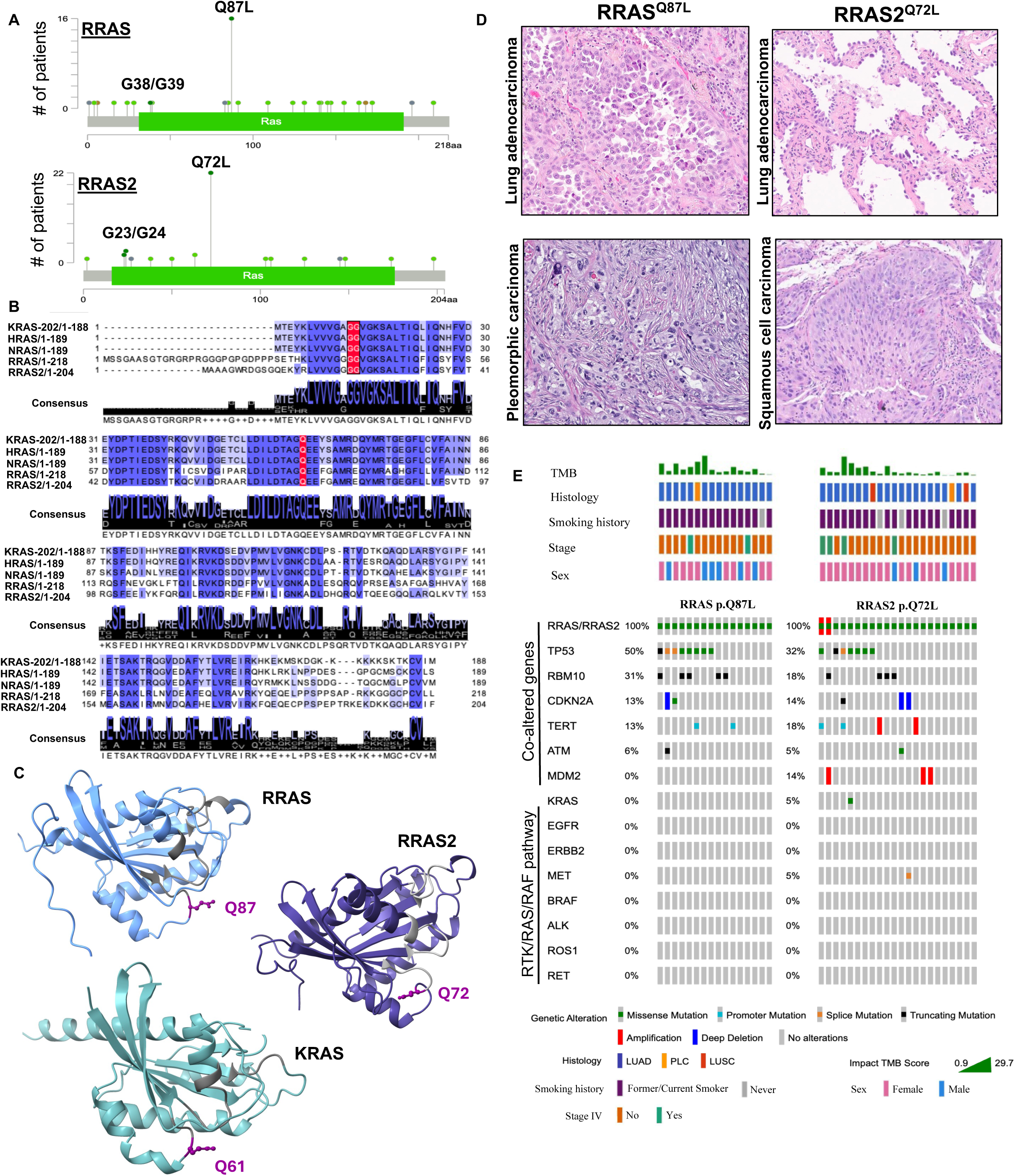
Characterization of *RRAS* and *RRAS2* mutations in lung cancer. **A**) Lollipop plots of *RRAS* (top panel) *or RRAS2* (bottom panel) mutations in NSCLC, demonstrating a clear hotspots for RRAS p.Q87L (top panel) and *RRAS2* p.Q72L. A single case of *RRAS2* p.Q72H was observed but not included in the figure. **B**) Primary amino acid sequence alignment of KRAS, NRAS, HRAS, RRAS, and RRAS2. Alignment of RAS family proteins performed using Jalview with Clustal Omega’s alignment algorithm^47^. **C**) 3D protein structure visualization of RRAS (AF-P10301-F1-v6), RRAS2 (AF-P62070-4-F1-v6), and KRAS (AF-P01116-F1-v6) captured from the AlphaFold Protein Structure Database^48,49^. GTP binding sites labeled in grey. **D)** Histological patterns of NSCLC with RRAS c.260A>T, p.Q87L (Left top: LUAD with lepidic pattern; Left bottom: pleomorphic carcinoma) or RRAS2 c.215A>T, p.Q72L mutations (Right top: LUAD with micropapillary pattern; Right bottom: LUSC). Images captured at 10x. **E**) Oncoprint of NSCLC with *RRAS* p.Q87L (left panel) and *RRAS2* p.Q72L (right panel). TMB measured in muts/Mb. aa=amino acid; RTK=receptor tyrosine kinase; TMB=tumor mutational burden; LUAD=lung adenocarcinoma; PLC=pleomorphic carcinoma of the lung; LUSC=lung squamous cell carcinoma.

### Clinical, pathologic, and genomic characteristics of *RRAS*^Q87L^ and *RRAS2*^Q72L^-mutated NSCLC

The median age at diagnosis for patients with *RRAS*^Q87L^-mutated lung cancers was 77.5 years, while those with *RRAS2*^Q72L^-mutated tumors was 70.5 years **(Table 1)**. The majority of patients with either mutation was female with a positive smoking history. The distribution of clinical stage at presentation was heterogenous and comparable to the *RRAS*/*RRAS2*-wild type LUAD comparison cohort. Two *RRAS*^Q87L^ mutations were found in lung adenocarcinoma *in situ* (AIS). Up to 32% and 45% of NSCLC with *RRAS*^Q87L^ and *RRAS2*^Q72L^ mutations, respectively, presented at stage III/IV. A review of tumor histology revealed that nearly all cases were comprised of lung adenocarcinomas (LUADs) (15/16, *RRAS*^Q87L^ and 19/22, *RRAS2*^Q72L^). The remaining patients had pleomorphic carcinoma of the lung (n=1, *RRAS*^Q87L^ and n=1, *RRAS2*^Q72L^) and lung squamous cell carcinoma (n=2, *RRAS2*^Q72L^) **(Figure 1D and Supplementary Figure 2)**.

**Table 1.** Clinical, pathologic and genomic characteristics of patients. *IASLC grades were assessed for resected invasive LUADs. PD-L1 positivity was defined as having a score of ≥1%. AIS and stage I have been combined for comparison of clinical stage. Abbreviations: AIS=adenocarcinoma *in situ*; LUAD=lung adenocarcinoma; LUSC=lung squamous cell carcinoma; IHC=immunohistochemistry; TMB=tumor mutational burden.

| Characteristics | RRAS p.Q87L<br>NSCLC<br>(n=16) |  | RRAS2 p.Q72L<br>NSCLC<br>(n=22) |  | RRAS/RRAS2<br>wild type<br>LUAD (n=520) |
| --- | --- | --- | --- | --- | --- |
| **Clinical data** |  | P value |  | P value |  |
| Median age, years (range) | 77.5 (54-86) | 0.00002 | 70.5 (61-81) | 0.04 | 67.2 (27.5-92.2) |
| Sex |  |  |  |  |  |
| Female | 10 (63%) | 1 | 19 (86%) | 0.04 | 336 (65%) |
| Male | 6 (38%) |  | 3 (14%) |  | 184 (35%) |
| Smoking |  |  |  |  |  |
| Ever | 15 (94%) | 0.05 | 19 (86%) | 0.2 | 318 (71%) |
| Never | 1 (6%) |  | 3 (14%) |  | 127 (29%) |
| Clinical stage |  |  |  |  |  |
| AIS | 2 (13%) | 0.2 | 0 (0%) | 0.2 | 0 (0%) |
| I | 9 (56%) |  | 10 (45%) |  | 275 (53%) |
| II | 0 (0%) |  | 2 (9%) |  | 61 (12%) |
| III | 3 (19%) |  | 6 (27%) |  | 58 (11%) |
| IV | 2 (13%) |  | 4 (18%) |  | 126 (24%) |
| **Pathologic data** |  | - |  | - |  |
| Histology |  |  |  |  |  |
| LUAD | 15 (94%) |  | 19 (86%) |  | 520 (100%) |
| LUSC | 0 (0%) |  | 2 (9%) |  | 0 (0%) |
| Pleomorphic | 1 (6%) |  | 1 (5%) |  | 0 (0%) |
| IASLC grade* |  | - |  | - |  |
| 1 | 0 (0%) |  | 4 (36%) |  | NA |
| 2 | 3 (50%) |  | 3 (27%) |  | NA |
| 3 | 3 (50%) |  | 4 (36%) |  | NA |
| **Biomarker data** |  |  |  |  |  |
| PD-L1 IHC |  |  |  |  |  |
| Positive | 4 (29%) | - | 9 (41%) | - | NA |
| Negative | 10 (71%) |  | 13 (59%) |  | NA |
| Median TMB, muts/Mb (range) | 7.2 (0.8-26.3) | 0.09 | 5.1 (0.9-29.7) | 0.9 | 5.2 (0-59.7) |

Focusing on LUAD, the most common histologic subtype, the combined frequency of *RRAS*^Q87L^ and *RRAS2*^Q72L^ mutations was 0.53% in LUAD patients screened for *RRAS* and *RRAS2* mutations (34/6,359). Among resected invasive LUADs, IASLC grade 3 was seen in 50% of *RRAS*^Q87L^-mutated cases and 36% of *RRAS2*^Q72L^-mutated cases. The majority of tumors were negative for PD-L1 expression by immunohistochemistry (71% and 59% PD-L1 negative among *RRAS*^Q87L^ and *RRAS2*^Q72L^ cases, respectively) and median tumor mutational burdens (TMB) for *RRAS*^Q87L^ and *RRAS2*^Q72L^ were 7.2 and 5.1 muts/Mb, respectively. Remarkably, all tumors with *RRAS*^Q87L^ lacked other canonical activating alterations in the receptor tyrosine kinase (RTK)/RAS/RAF pathway. Similarly, tumors with *RRAS2*^Q72L^ were also wild type for other activating RTK/RAS/RAF alterations, with the exception of two cases that had concurrent MAPK pathway mutations: one KRAS^G12V^ and one MET exon 14 skipping mutation. Other recurrently co-altered genes included *TP53*, *RBM10*, *CDKN2A*, *TERT* and *ATM* **(Figure 1E)**.

Next, we explored whether the *RRAS*^Q87L^ and *RRAS2*^Q72L^ mutations were clonal by using different methods to infer clonality. In samples with FACETS data available, we calculated cancer cell fraction (CCF) for *RRAS*^Q87L^ and *RRAS2*^Q72L^ mutations, using >80% as the cutoff for calling mutations clonal. Out of 10 samples where FACETS data were available and evaluable (three *RRAS*^Q87L^ and seven *RRAS2*^Q72L^), all 10 cases showed CCF that was >80%, supporting the clonal nature of *RRAS*^Q87L^ and *RRAS2*^Q72L^ mutations. In addition, two samples with *RRAS2*^Q72L^ had amplification of *RRAS2* mutant allele as reflected by variant allele fractions (VAFs) of the *RRAS2*^Q72L^ of 88% and 92%, consistent with clonality. In the remaining samples without FACETS data (n=26), we inferred clonality based on the relative VAF fraction (RVF) of each mutation. Similar to CCF, we categorized mutations with >80% RVF as likely clonal. Three cases lacked other non-synonymous mutations besides the hotspot mutations in *RRAS*/*RRAS2*, establishing these variants in the three cases as de facto clonal drivers. One case had only one mutation (*SPEN* p.S3389Ffs*26) besides *RRAS2*^Q72L^ and was therefore considered not amenable to the RVF analysis. In the remaining 22 samples, all but one had an RVF that was >80% (12 *RRAS*^Q87L^ and 9 *RRAS2*^Q72L^), consistent with clonality in 21 of these 22 cases. One sample with *RRAS2*^Q72L^ had an RVF of 63%. Interestingly, subsequent molecular testing of a different block from the same tumor demonstrated a similar genomic profile but without the *RRAS2*^Q72L^ mutation, supporting its subclonal nature in this single case.

To assess the clinico-genomic features associated with *RRAS*^Q87L^ and *RRAS2*^Q72L^ mutations, we compared our cohorts with a stage-matched control group of LUADs lacking *RRAS*^Q87L^ and *RRAS2*^Q72L^ mutations. The clinical and molecular characteristics of this cohort have been published previously.^34^ Comparative analysis revealed that patients with *RRAS*^Q87L^ and *RRAS2*^Q72L^ were older than patients with no mutations in these two genes. In addition, patients with *RRAS2*^Q72L^ mutations were more likely to be women compared to the control group. There were no other significant differences in clinical, pathologic, and genomic features, including co-mutational landscapes. In terms of their clinical outcomes, the median overall survival (OS) for patients with *RRAS*^Q87L^ and *RRAS2*^Q72L^-mutated NSCLC were 44 months (95% CI: 29-not estimable) and 85 months (95% CI: 20-not estimable), respectively. There was no significant difference in the median OS between the two cohorts (logrank test, p=0.6) (**Supplementary Figure 3**). The full demographic, clinical and genomic characteristics of patients with *RRAS*^Q87L^ and *RRAS2*^Q72L^ are provided in **Supplementary Table 4A and 4B**.

### *RRAS* and *RRAS2* mutations in other large WES cohorts

To further explore the characteristics of *RRAS* and *RRAS2* mutations and to correlate our findings with those from other independent cohorts, we searched for *RRAS* and *RRAS2* mutations in the TCGA LUAD (n=566) and LUSC (n=484) whole-exome sequencing (WES) data sets. In total, three *RRAS* and five *RRAS2* mutations were identified. *RRAS*^Q87L^ and *RRAS2*^Q72L^ were the most common and only recurrent mutations, with each variant seen in two cases. The other remaining mutations were observed once and were all classified as variants of unknown significance. Notably, mutations involving p.G38/G39 in *RRAS* and p.G23/G24 in *RRAS2* (corresponding to p.G12/G13 in classical RAS) were not seen in those lung cancer cohorts. In terms of histology, both mutations were present equally in LUAD and LUSC. Similar to our MSK cases, all tumors with *RRAS*^Q87L^ and *RRAS2*^Q72L^ lacked other activating alterations in the RTK/RAS/RAF pathway. In addition to the TCGA cohorts, we searched for *RRAS* and *RRAS2* mutations in another large WES LUAD series (OncoSG, n=302). In this series, *RRAS2*^Q72L^ was identified in a single tumor that was also wild type for activating alterations in other RTK/RAS/RAF genes. Taken together, the overall frequency of *RRAS*^Q87L^ and *RRAS2*^Q72L^ was 0.37% within NSCLC cases (5/1,352) and 0.35% within LUAD (3/868), from the respective WES cohorts. This was largely concordant with the frequencies observed in our MSK series. The full clinical and pathologic features of *RRAS*^Q87L^- and *RRAS2*^Q72L^-mutated NSCLC cases from other WES cohorts are provided in **Supplemental Table 4C**.

### RRAS^Q87L^ and RRAS2^Q72L^ are oncogenic and targetable mutations in Ba/F3 cells

To investigate the oncogenic potential of these clinically identified *RRAS* and *RRAS2* mutations *in vitro*, we first expressed the mutations of interest in the murine pro-B cell line Ba/F3. Ba/F3 cells are a popular model for studying oncogenesis due to their dependence on interleukin-3 (IL-3) for proliferation and loss of this dependency when cells are transformed.^35^ Expression of RRAS^Q87L^ or RRAS2^Q72L^ in Ba/F3 cells resulted in IL-3-independent growth while growth of cells expressing an empty vector or wild-type RRAS or RRAS2 was halted when IL-3 was withdrawn **(Figure 2A)**. Ba/F3 cells expressing RRAS^Q87L^ or RRAS2^Q72L^ also formed tumors in NSG mice **(Supplementary Figure 4)**. Increased phosphorylation of Erk 1/2 and Akt was observed in Ba/F3 cells expressing RRAS^Q87L^ or RRAS2^Q72L^ compared to cells expressing the wildtype counterparts **(Figure 2B**). The RRAS2 antibody (Proteintech 12530-1-AP) demonstrated some cross-reactivity with RRAS at all attempted antibody concentrations and blot exposure times.

**Figure 2.**
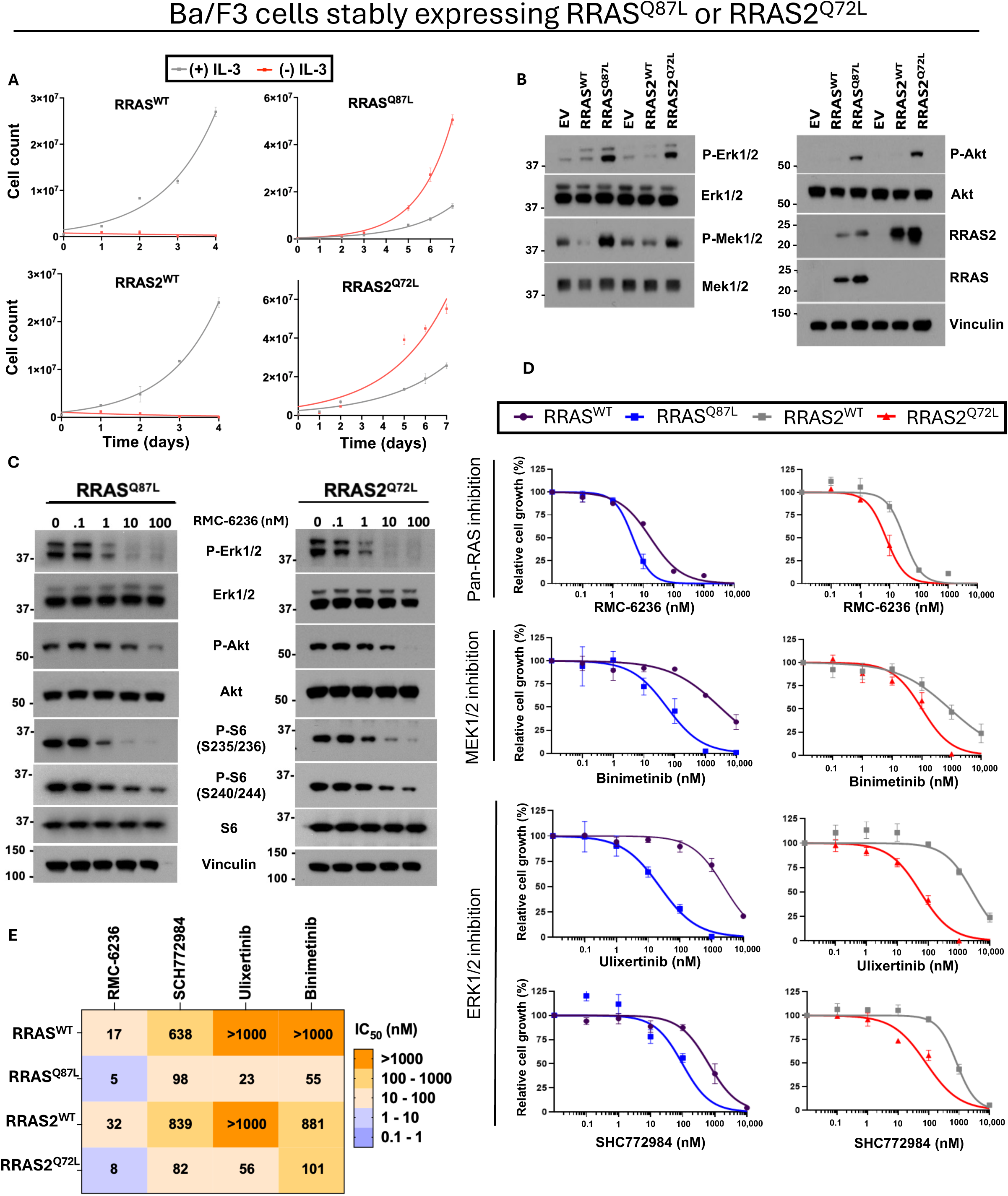
Expression of RRAS^Q87L^ or RRAS2^Q72L^ is oncogenic in Ba/F3 cells. **A**) Growth curves of Ba/F3 cells expressing an empty vector (EV), wild-type (WT) RRAS/RRAS2, or mutant RRAS/RRAS2 grown in media with or without 5 ng/mL IL-3. **B-C**) Whole-cell extracts from isogenic Ba/F3 cells left untreated or treated with the indicated RMC-6236 concentrations were resolved by SDS-PAGE and immunoblotted for the indicated proteins. Representative immunoblots from two independent experiments are shown with quantitation shown in **Supplementary Figure 6. D**) The indicated Ba/F3 cell lines were treated for 96 h with inhibitors of ERK 1/2 (SHC772984, ulixertinib), MEK1 /2 (binimetinib), or RAS (RMC-6236). Results represent the mean ± SEM of three independent experiments where each condition was assayed in triplicates. **E)** Heat map of IC_50_ values. IC_50_ values with 95% confidence intervals for growth inhibition are given in **Supplemental Table 5.**

To compare the signaling properties of RRAS^Q87L^ and RRAS2^Q72L^ mutations with other RAS hotspots that were not enriched in the clinical sequencing data, HEK-293T cells were transiently transfected to express the K/N/HRAS G12 homologous mutations RRAS^G38C/D^ and RRAS2^G23C/D^ in addition to RRAS^Q87L^ and RRAS2^Q72L^. Expression of RRAS^Q87L^ and RRAS2^Q72L^ led to increased MAPK pathway activation relative to expression of the wildtype counterparts as observed in Ba/F3 cells. RRAS2^G23D^ led to increased ERK 1/2 and P90 RSK phosphorylation, while RRAS^G38C/D^ and RRAS2^G23C^ resulted in no observed changes to ERK 1/2, P90 RSK, or S6 phosphorylation. All RRAS and RRAS2 mutants led to some degree of increased AKT phosphorylation, though the effects were subtle relative to MAPK activation **(Supplementary Figure 5)**.

We next sought to identify therapeutic vulnerabilities of Ba/F3 cells expressing RRAS^Q87L^ or RRAS2^Q72L^ utilizing MAPK-targeted therapies including RMC-6236. RMC-6236 is a first-in-class, multi-selective inhibitor of GTP-bound RAS targeting the active state of multiple members of the RAS family, including both wild-type and mutant variants.^10,36^ The preclinical efficacy of RMC-6236 has been demonstrated in cells harboring mutant KRAS, NRAS, and HRAS with mutations at codons 12, 13, and 61. We thus set out to determine if RMC-6236 also possesses activity against RRAS and RRAS2 mutants to potentially expand the use of this class of inhibitors to additional members of the RAS family GTPases. RMC-6236 is currently under investigation in multiple clinical trials (NCT05379985, NCT06625320, NCT06040541; NCT06162221; NCT06445062; NCT06128551).

Western blotting analysis of cell extracts prepared from Ba/F3 cells expressing either wild-type or mutant RRAS/RRAS2 indicated that RRAS-and RRAS2-mutant cells treated with RMC-6236 exhibited diminished phosphorylation of Erk 1/2, Akt and S6 in a dose-dependent manner, with greater sensitivity seen in Mapk signaling compared to Akt (**Figure 2C**). The capacity of RMC-6236 to prevent activation of members of both the Mapk and Pi3k-Akt-mTor pathway has been demonstrated in cell models of KRAS^G12X^.^36^ RMC-6236 led to decreased Erk1/2 phosphorylation in RRAS^Q87L^-and RRAS2^Q72L^-mutant Ba/F3s at 1-100 nM. The RRAS^WT^-Ba/F3s appeared to exhibit a small but insignificant degree of re-activation of Erk 1/2 phosphorylation at 100 nM. RMC-6236 treatment led to reduced Erk 1/2 phosphorylation in RRAS2^WT^ cells at 1-100 nM though to a lesser extent than observed in Ba/F3-RRAS2^Q72L^ cells **(Figure 2C and Supplementary Figure 6)**. In concordance with this data, RRAS^Q87L^ and RRAS2^Q72L^ mutants demonstrated enhanced sensitivity to targeted inhibitors of MEK1/2 (binimetinib), ERK1/2 (SHC772984, ulixertinib), and RAS (RMC-6236), relative to Ba/F3 cells with overexpression of wild-type RRAS or RRAS2 (**Figure 2D-E and Supplementary Table 5**).

### Human lung cells expressing RRAS^Q87L^ or RRAS2^Q72L^ are transformed and sensitive to RMC-6236

We next set out to investigate the effect of overexpression of RRAS^Q87L^ or RRAS2^Q72L^ in an immortalized non-transformed human lung cell line [human bronchiolar epithelial cells (HBEC)]. RRAS^Q87L^ and RRAS2^Q72L^ were again sufficient to transform HBEC cells *in vivo*, with tumor formation following subcutaneous injection into NSG mice **(Figure 3A)**. HBEC-RRAS2^Q72L^ xenograft tumors grew faster than HBEC-RRAS^Q87L^. Similar to Ba/F3 cells, HBEC cells overexpressing the *RRAS* and *RRAS2* mutants had enhanced activation of ERK1/2 and downstream signaling proteins such as S6 and P90 RSK, as well as activation of AKT, P70 RSK, and STAT3 **(Figure 3B)**. HBEC cells expressing mutant RRAS/RRAS2 showed upregulation of negative regulators of the MAPK pathway, including SPRED1, SPRY1, and SPRY2 **(Figure 3B)**. Increased expression of DUSP6 was more prominent in cells expressing RRAS2^Q72L^ compared to cells expressing wild-type RRAS/RRAS2 or RRAS^Q87L^ **(Figure 3B)**. It is possible that the negative regulators of the MAPK pathway were upregulated to restrain activation of MEK 1/2 and ERK 1/2, thereby preventing cell death or senescence due to ERK 1/2 hyperactivation. This phenomenon has been observed in other models of RAS-driven cancer, including pancreatic and colorectal cancers.^37^

**Figure 3.**
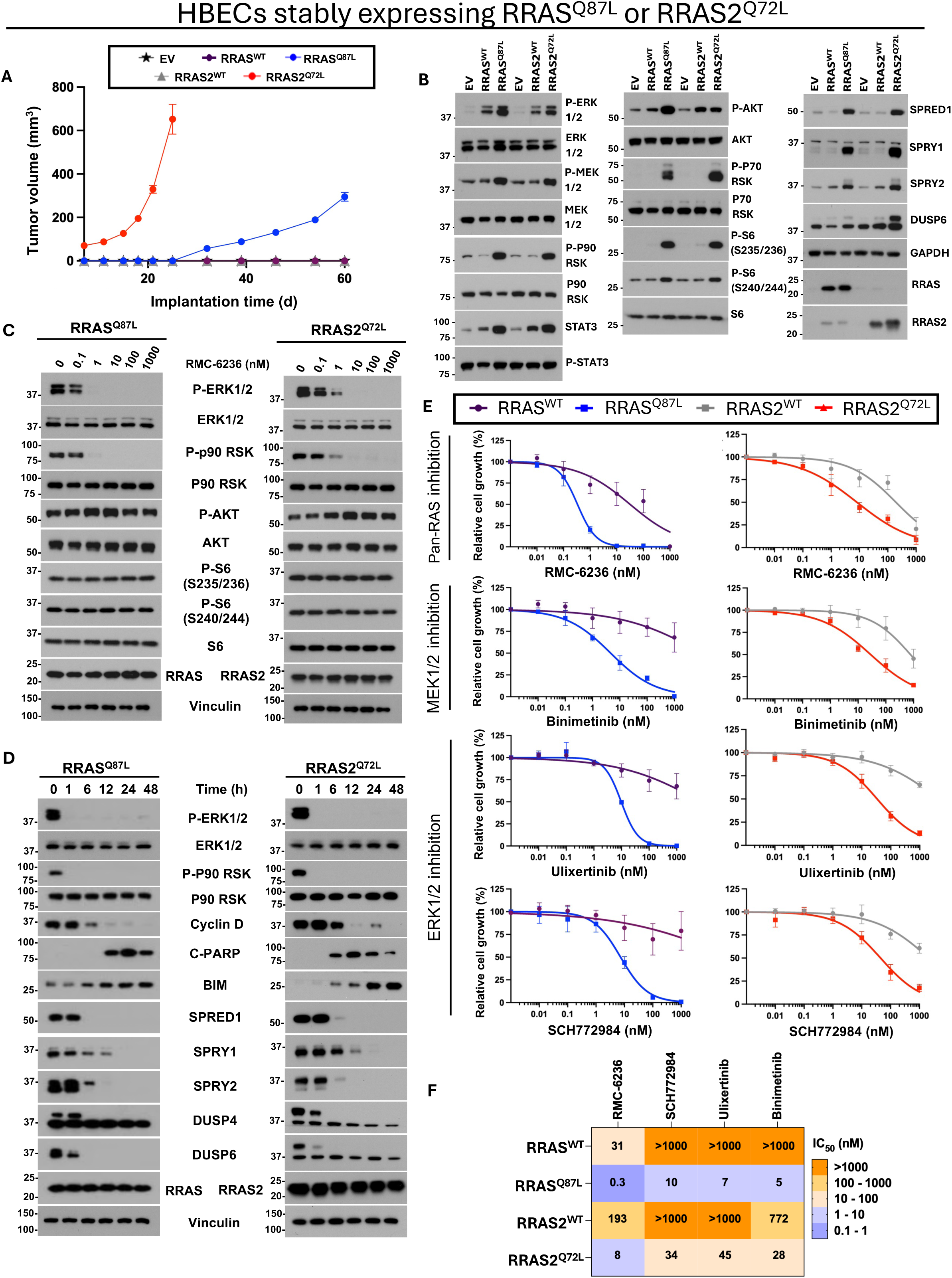
Human bronchiolar epithelial cells expressing RRAS^Q87L^ or RRAS2^Q72L^ are oncogenic and sensitive to pan-RAS inhibition. **A**) Tumor growth in NSG mice following subcutaneous injection of HBEC cells expressing an empty vector (EV), RRAS/RRAS2 wild type (WT), or mutant RRAS/RRAS2. Data represent the mean ± SEM of six tumors. **B-D**) Whole-cell extracts were subjected to Western blot analysis. Cells were left untreated (**B**), treated for 90 mins with the indicated concentrations of RMC-6236 (**C**), or treated with 10 nM RMC-6236 for the time shown (**D**). Representative immunoblots from two independent experiments are shown. **E)** Isogenic HBEC cells were treated for 96 h with the indicated concentrations of inhibitors targeting ERK 1/2 (SHC772984, ulixertinib), MEK 1/2 (binimetinib), or RAS (RMC-6236). Results represent the mean ± SD of 2 independent experiments in which there were three replicates of each condition. **F)** Heat map of IC_50_ values. The IC_50_ values with 95% confidence intervals for growth inhibition are given in **Supplemental Table 5.**

RRAS^Q87L^ and RRAS2^Q72L^ mutant HBECs treated with RMC-6236 demonstrated diminished phosphorylation of ERK 1/2 and P90 RSK in a dose-dependent fashion, though phosphorylation of AKT and the downstream effector S6 showed no reduction **(Figure 3C)**. Treatment with 10 nM RMC-6236 over a 48-hour time course also resulted in sustained inhibition of ERK 1/2 and P90 RSK phosphorylation, downregulation of negative regulators of the MAPK pathway and markers of cell cycle progression, and upregulation of pro-apoptotic proteins **(Figure 3D).** HBEC cells harboring RRAS^Q87L^ or RRAS^Q87L^ exhibited enhanced sensitivity to inhibitors of the MAPK pathway relative to cells expressing the wild-type counterparts **(Figure 3E-F and Supplemental Table 5)**.

To investigate *in vivo* efficacy of RMC-6236, NSG mice bearing HBEC-RRAS^Q87L^ or HBEC-RRAS2^Q72L^ xenograft tumors were treated with vehicle or 10 mg/kg-50 mg/kg RMC-6236 once daily (**Figure 4A-B)**. Histology with mitotic index (bottom panels), tumor volume over time (left panels), area under curve analysis (middle panels), and percentage tumor volume change from baseline (right panels) are shown. In both xenograft models RMC-6236 caused a significant reduction in tumor growth compared to vehicle-treated animals. The dose of 25 mg/kg was as effective as 50 mg/kg in RRAS2^Q72L^ tumors. The 50 mg/kg dose was more effective in the HBEC-RRAS^Q87L^ xenograft model, leading to nearly complete tumor stasis. Nevertheless, the 25 mg/kg and 50 mg/kg doses caused similar reductions in the number of mitotic cells seen on microscopy (Figure 4A-B, bottom, left panels). Histologic examination showed that untreated mice grew high grade tumors, as seen by solid growth, abundant pleomorphism and higher rates of cell division. Notably, treated tumors developed squamous and mucinous features, suggesting partial differentiation to native bronchial epithelium **(Figure 4A-B, bottom panels)**. To assess target engagement, we performed Western blotting analysis on HBEC-RRAS2^Q72L^ xenograft tumors excised at the end of the study. Tumors treated with vehicle or 25 mg/kg RMC-6236 (three tumors per group) showed a significant reduction in phosphorylation of S6 at both the S240/244 (canonical marker of mTOR activity^38^) and S235/236 (target of P90 RSK downstream of ERK 1/2, as well as mTOR ^39^) sites. Although ERK 1/2 phosphorylation was diminished by 58.3 ± 3.4% this was not statistically significant (p=0.055) **(Figure 4C)**. Tumors treated with RMC-6236 showed significant induction of BIM **(Figure 4C)**. Importantly, AKT-S473 phosphorylation was not affected by RMC-6236 treatment **(Figure 4C)**. Tumors from the HBEC-RRAS^Q87L^ xenograft model were not available to perform similar Western blotting analysis. No reduction in animal weight **(Supplementary Figure 7)** or dose-dependent toxicity was observed up to 50 mg/kg.

**Figure 4.**
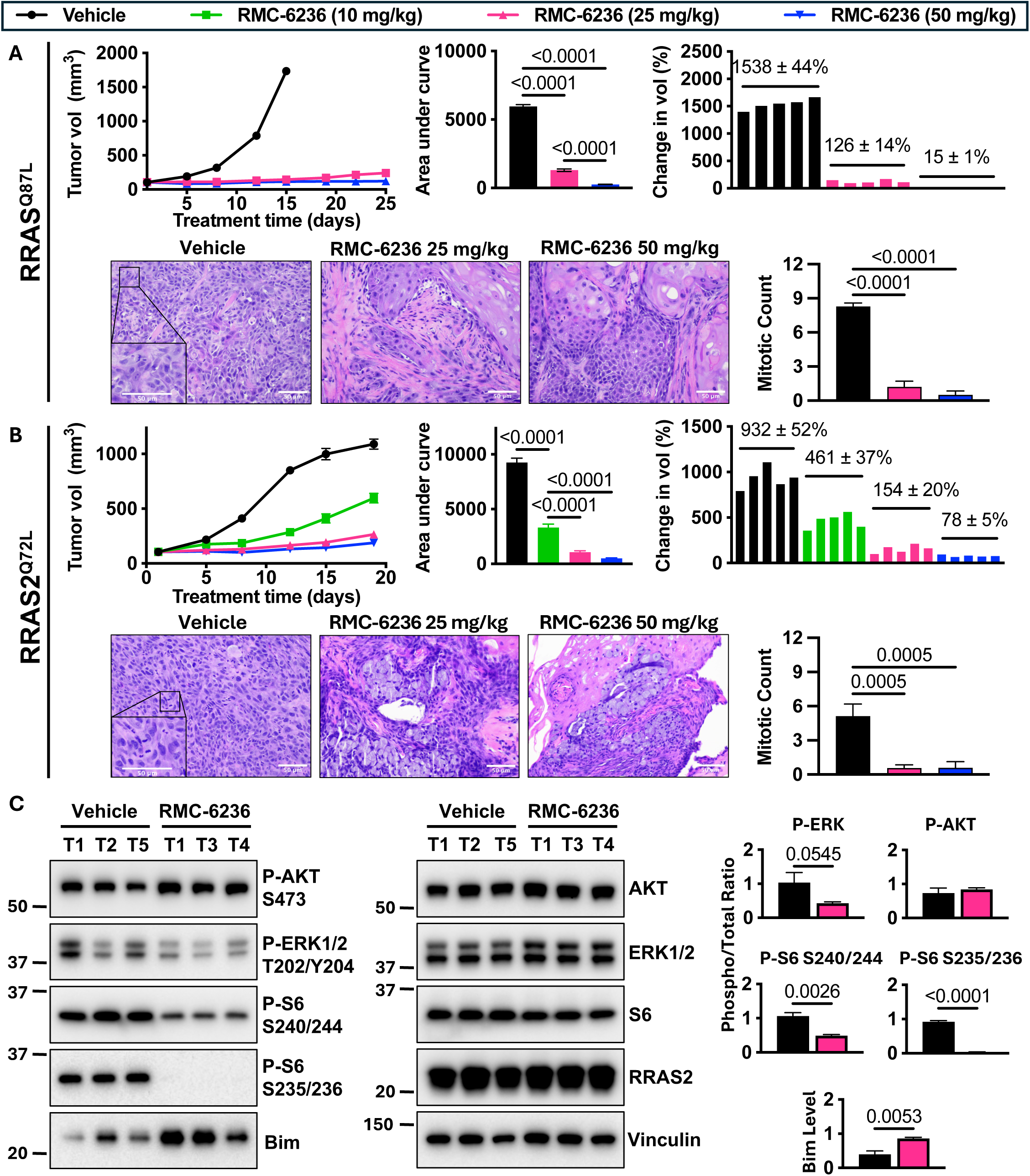
RMC-6236 is effective therapy in *in vivo* models of RRAS^Q87L^ or RRAS2^Q72L^ driven lung cancer. **A-B**) Tumor growth in immunocompromised NSG mice treated with RMC-6236 (vehicle, 10 mg/kg, 25 mg/kg, or 50 mg/kg, once daily (q.d.)). Top left: tumor volume measurements; Top middle: area under curve (AUC) analysis of tumor volume over the treatment course; Top right: tumor volume change from baseline (expressed as %) of individual tumors with the average ± SEM of the group displayed middle. All results represent the mean ± SEM (n = 5). Measurements were compared with one-way ANOVA with Tukey’s multiple comparisons test. AUC values for each treatment groups compared to vehicle, p < 0.0001. Representative H&E staining for the indicated treatment condition (scale bar represent 50 um) and inset demonstrating mitotic figures for vehicle treatment are shown in the bottom panels. Average number of mitotic counts per high-powered field (40X objective) encompassing 10 fields per tumor averaged over three tumors per RRAS/RRAS2 mutant model is shown in the graph on the right. **C**) Western blot analysis of three xenograft tumors from RRAS^Q72L^-tumor bearing mice treated with vehicle (V1, V2, V5) or 25 mg/kg RMC-6236 daily (T1, T3, T4). Phospho/total protein ratios are shown. Data represents the mean of three tumors ± SEM. Comparisons made with unpaired t tests.

## DISCUSSION

The present findings nominate RRAS^Q87L^ and RRAS2^Q72L^ as novel and recurrent oncogenic drivers in lung cancers, found in approximately 0.37-0.45% of NSCLC samples in the MSK-IMPACT cohort and other large WES NSCLC series. The observed prevalence of RRAS^Q87L^ and RRAS2^Q72L^ is similar to other well-studied and clinically relevant driver oncogenes in NSCLC including HRAS, NRAS, ROS1, NTRK, and NRG1, all present at approximately 0.1-1% frequency.^40^ Importantly, RRAS^Q87L^ or RRAS2^Q72L^ mutations were almost entirely non-overlapping with other activating RTK-RAS-RAF alterations and predominantly clonal, suggesting that these mutations function as a primary oncogenic driver. A smaller number of *RRAS/RRAS2* mutations homologous to other KRAS hotspots such as G12/G13 (RRAS^G38/G39^ and RRAS2^G23/G24^) were also identified in this study but these mutations were not highly recurrent.

It remains uncertain as to why these latter mutations are not enriched in RRAS-mutant LUADs despite their predominance in other RAS-mutant cancers. Mutation-specific differences in cell signaling and tumor type prevalence is a well-appreciated aspect of RAS biology in cancer.^41^ In HEK-293T cells, RRAS^G38C/D^ and RRAS2^G23C^ did not lead to enhanced phosphorylation of ERK, AKT, or the downstream effectors P90 RSK and S6 to the same degree as RRAS^Q87L^ or RRAS2^Q72L^. Hence, it appears KRAS G12-homologous mutations in RRAS/RRAS2 are not sufficient to drive MAPK hyperactivation in human cells, and their ability to drive tumorigenesis may be dependent on other, co-existing driver mutations.

Expression of RRAS^Q87L^ or RRAS2^Q72L^ alone were sufficient to transform murine Ba/F3 cells and human lung cells. RRAS2^Q72L^ has previously been identified as capable of forming tumors *in vivo*,^42^ though our study is the first direct demonstration of the oncogenic role of RRAS^Q87L^. These mutations resulted in activation of both the MAPK and AKT-PI3K-mTOR signaling axes, as well as potently upregulating negative regulators of MAPK function. Increased expression of these negative regulators may be necessary to prevent hyperactivation of ERK 1/2, which can be toxic to cells. Fernandez-Pisonero et al. previously demonstrated somatic induction of RRAS2^Q72L^ in mice primarily drives PI3K and/or mTOR dependent tumors including lymphoblastic leukemias and ovarian cystadenomas.^42^ Our results suggest RRAS^Q87L^ and RRAS2^Q72L^ mutations are addicted to the MAPK pathway as inhibitors of MEK 1/2 or ERK 1/2 block growth of cell lines. In RRAS^Q87L^ and RRAS2^Q72L^-mutant lung cells, phosphorylation of ERK 1/2 and its downstream effector P90 RSK was potently inhibited by the pan-RAS inhibitor RMC-6236 at similar concentrations required in KRAS^G12^-mutant models. *In vivo*, RAS inhibition with RMC-6236 inhibited tumor growth and reduced MAPK pathway activation without any appreciable reduction in AKT phosphorylation. Although our study indicates that RRAS^Q87L^ and RRAS2^Q72L^ mutations appear to drive tumorigenesis primarily through the MAPK pathway in human lung cells, and this dependency is targetable with RMC-6236, we are unsure of the role played by PI3K-AKT. Of note, we did not observe any tumor regression, despite seeing activation of pro-apoptotic pathway. It is possible that inhibition of the PI3K-AKT-mTOR pathway in combination with RRAS/RRAS2 inhibition may be necessary to induce sustained tumor regression.

Though targeted therapies against members of the MAPK pathway often show excellent preclinical efficacy in RAS-driven models of lung cancers, the clinical efficacy of these inhibitors is often hampered by drug resistance including on-target mutations or re-wiring of growth factor signaling.^43,44^ In an effort to overcome common drug resistance mechanisms as well as target multiple RAS isoforms, RMC-6236 was developed as a first-in-class, non-covalent inhibitor of GTP-bound RAS. RMC-6236 has shown efficacy in preclinical models with K/H/N-RAS, including mutations at the Q61 position and in settings of resistance to other RAS inhibitors.^36^ RMC-6236 is currently under investigation in multiple clinical trials in patients harboring RAS mutant solid tumors including NSCLC, pancreatic, and GI cancers. Available data from an ongoing phase 1/1b clinical trial (NCT05379985) show promising efficacy in previously treated, RAS-mutant pancreatic adenocarcinoma patients, with median progression-free survival of 8.5 months and 7.6 months in KRAS G12X and other RAS mutant tumors, respectively.^45^

In HBEC-RRAS^Q87L^ and HBEC-RRAS2^Q72L^ xenograft tumors, RMC-6236 inhibited tumor growth in a dose-dependent fashion (10 mg/kg – 50 mg/kg). Although dose titration to maximize clinical efficacy with a favorable tolerability profile is still ongoing, current trials of RMC-6236 monotherapy are utilizing daily dosing between 120-300 mg. Previous pharmacokinetic modeling has estimated the 300 mg daily dosing in humans would be needed to reach an equivalent mean blood exposure observed in mice treated with the 25 mg/kg daily dosing schedule.^36^ Hence, RMC-6236 appears to be active in RRAS and RRAS2-mutant cells at clinically achievable concentrations, though whether differences in activity compared to KRAS-mutant models is clinically significant remains to be determined. Of note, the pan-RAS inhibitors RMC-6236 and RMC-7977 have demonstrated greater efficacy against RAS G12 mutants versus G13 or Q61, possibly owing to restoration of GTP hydrolysis in the former.^46^ It remains to be seen in clinical trials if non-G12 mutations retain adequate sensitivity at doses that do not exceed tolerability. For all RAS-mutant tumors, it is likely that combination therapy approaches will be important for maximal depth of response and durable clinical benefit. Multiple clinical trials are evaluating the potential of this approach, combining pan-RAS inhibitors with other targeted therapies, anti-PD1 blockade, and chemotherapy (NCT 06445062, NCT06162221, NCT06922591). Nevertheless, the similar signaling profiles and drug sensitivities observed in RRAS and RRAS2 mutant models relative to KRAS warrant inclusion of these mutations in the broader study of RAS-driven cancers.

The introduction of targeted therapies as front-line management of oncogene-driven advanced NSCLCs has revolutionized lung cancer treatment in the past two decades. However, there still remains approximately 20-30% of NSCLC tumors that lack a routinely identified, actionable oncogenic driver.^40^ Hence, there exists a pressing need to continue identifying actionable driver alterations in order for these patients to reap the benefits of existing and novel targeted therapies. By leveraging a comprehensive NGS panel that is designed for both clinical actionability and biomarker discovery, we have identified, for the first time, RRAS^Q87L^ and RRAS2^Q72L^ as recurrent mutations in driver-negative NSCLCs. Importantly, our experimental and functional data indicate that these mutations are oncogenic drivers vulnerable to the novel and clinically active pan-RAS inhibitor RMC-6236. Taken together, our findings delineate a novel cohort of patients with RRAS^Q87L^- and RRAS2^Q72L^-mutant lung cancers who may be potential candidates for future trials of RAS-targeted therapies. Notably, given that *RRAS* and *RRAS2* are not profiled on most NGS panels, addition of these genes in clinical sequencing panels should be considered in order to expand screening and to advance treatment strategies for patients with RRAS^Q87L^ and RRAS2^Q72L^-mutated lung cancers.

## ACKNOWLEDGEMENTS

This research was partly supported by a developmental research grant from the Department of Pathology and Laboratory Medicine at MSK (to S-RY) and grants from the National Cancer Institute of the National Institutes of Health, including P30 CA008748 (Cancer Center Support grant to MSK), 5R25CA020449 (Medical Summer Student Research Support for AJP, TZ, RC, RPM), and 5P01CA129243-13 (ML). The authors thank Dr. Monika Davare for her help with figure visualization. The authors also thank Tracey Riano for administrative help with the Medical Summer Student Research Program.

