## Supplementary Figures for "*RRAS* and *RRAS2* mutations are recurrent oncogenic drivers in lung cancer and are sensitive to the pan-RAS inhibitor RMC-6236"

### Supplementary Figure 1

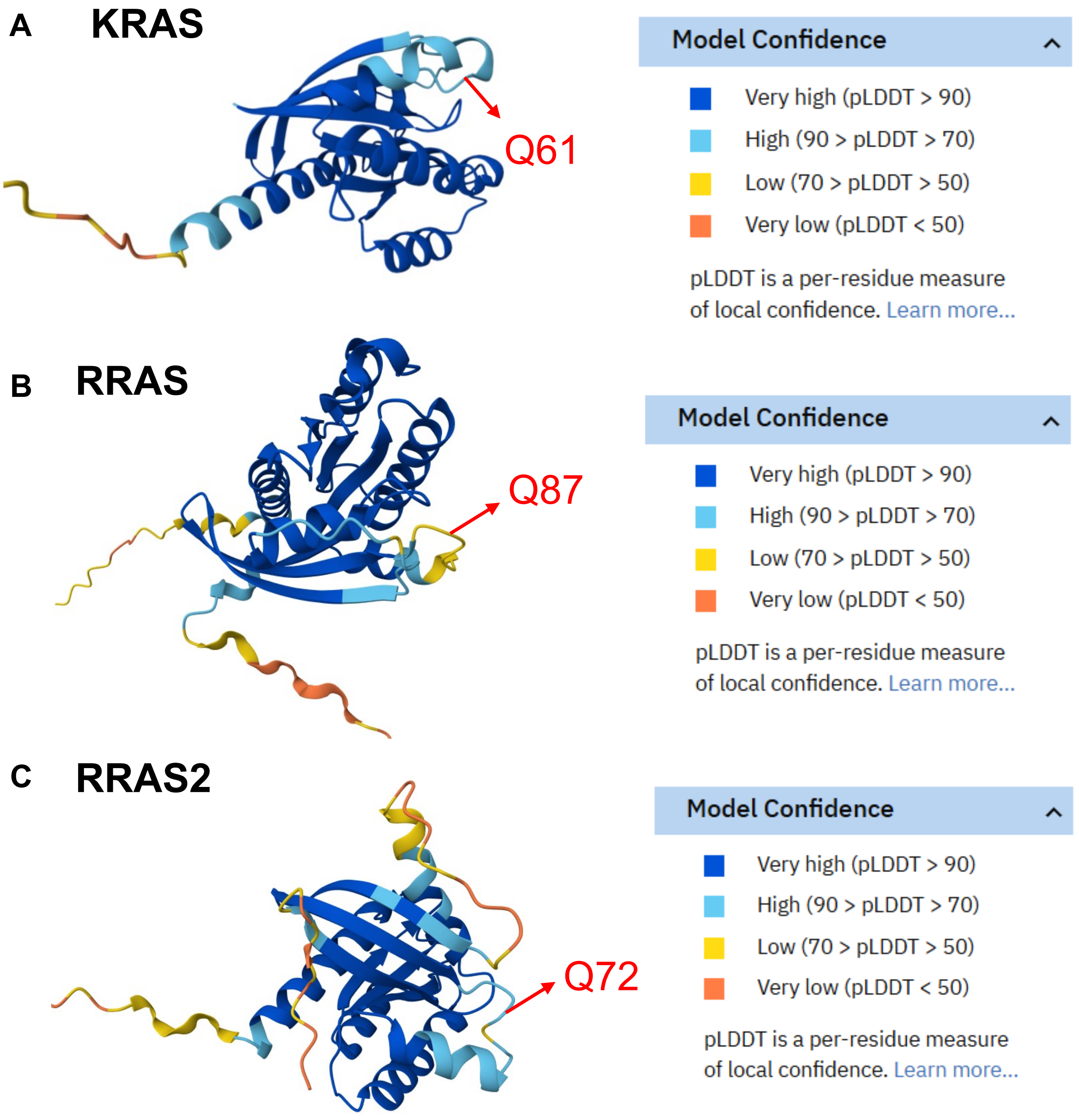


**Supplementary Figure 1. (A-C).** Protein structure visualization of RRAS (AF-P10301-F1-v6), RRAS2 (AF-P62070-4-F1-v6), and KRAS (AF-P01116-F1-v6) (left) was obtained from the AlphaFold Protein Structure Database. Model confidence of in silico structure prediction for loop structure containing mutations of interest (right).

### Supplementary Figure 2

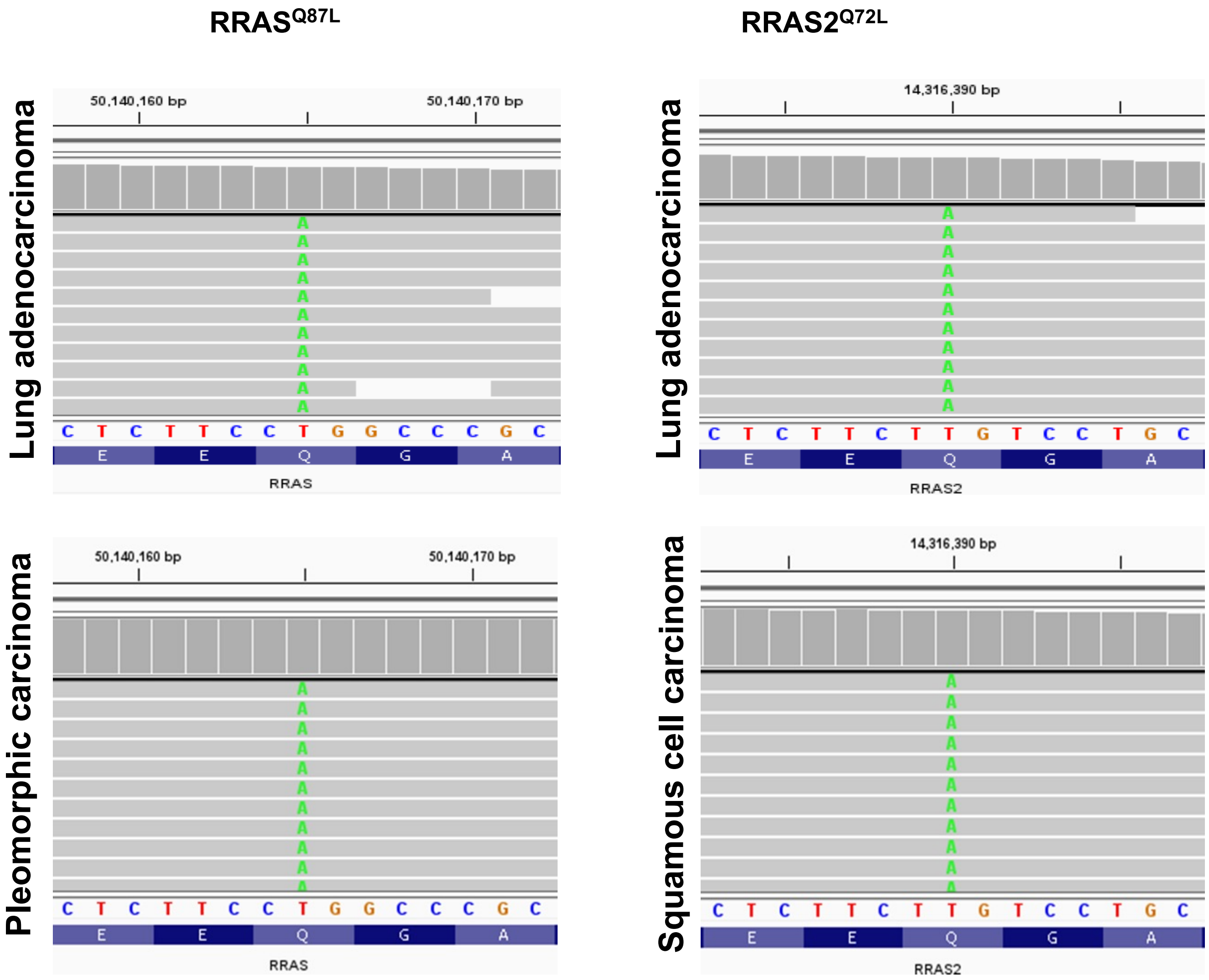


**Supplementary Figure 2.** Tumor genomic sequencing snapshot (IGV) for cases outlined in **Figure 1D**, including lung adenocarcinoma with micropapillary pattern and an RRAS c.260A>T, p.Q87L mutation, lung adenocarcinoma with lepidic pattern and an RRAS2 c.215A>T, p.Q72L mutation, pleomorphic carcinoma of the lung with an RRAS c.260A>T, p.Q87L mutation, lung squamous cell carcinoma with an RRAS2 c.215A>T, p.Q72L mutation.

### Supplementary Figure 3

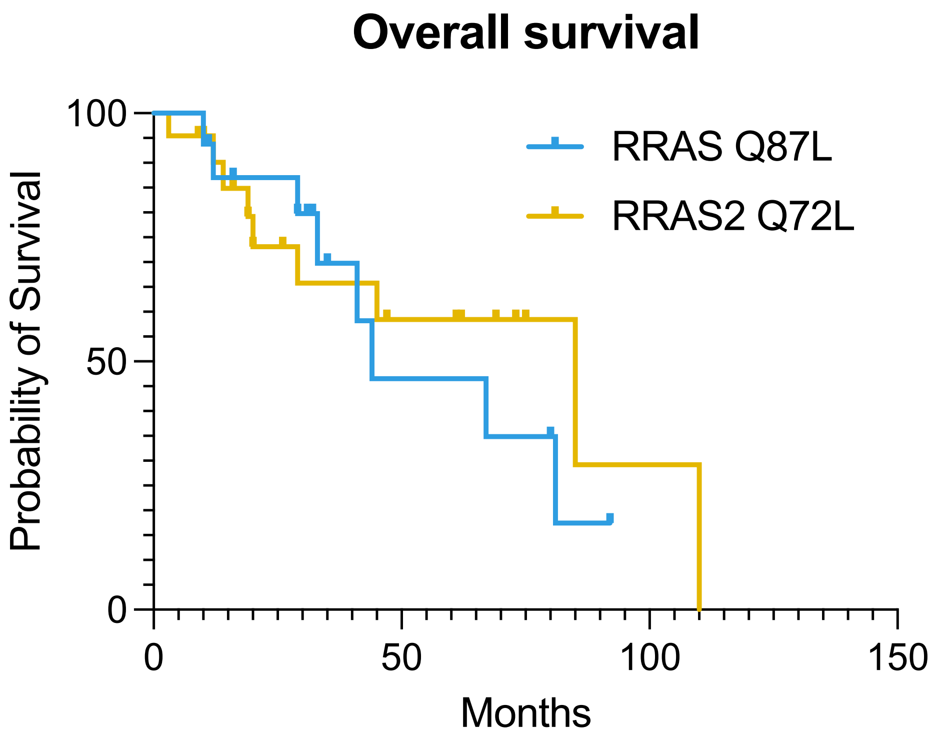


**Supplementary Figure 3.** Median overall survival (OS) for patients with *RRAS*$^{Q87L}$ and *RRAS2*$^{Q72L}$-mutated NSCLC were 44 months (95% CI: 29-not estimable) and 85 months (95% CI: 20-not estimable), respectively. There was no significant difference in the median OS between the two cohorts (logrank test, p=0.6).

### Supplementary Figure 4

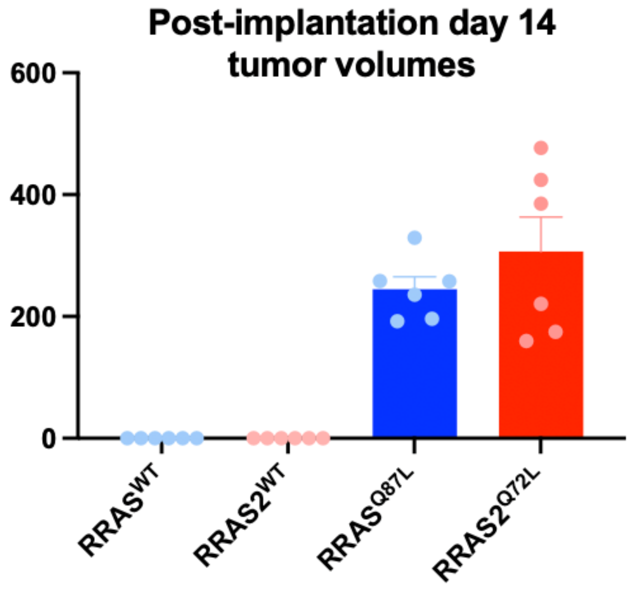


**Supplementary Figure 4.** Ba/F3 cells stably expressing RRAS$^{WT}$, RRAS2$^{WT}$, RRAS$^{Q87L}$, or RRAS2$^{Q72L}$ were implanted into the subcutaneous flank of NSG mice. Tumor volume at 14 days post-implantation is shown. Data represents mean $\pm$ SEM of 4 tumors in independent animals.

### Supplementary Figure 5

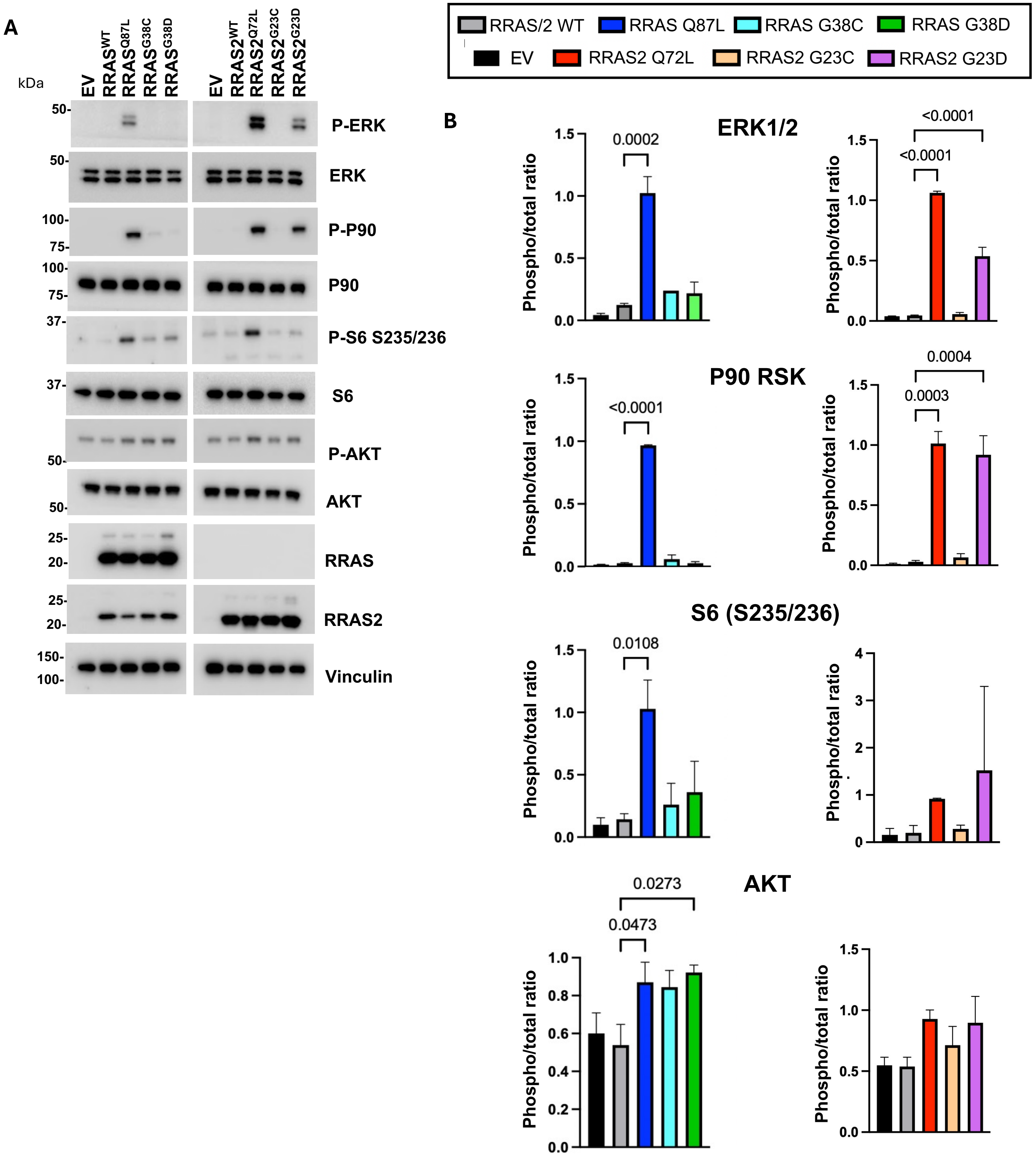


**Supplementary Figure 5.** Western blot analysis of HEK-293T cells transiently transfected with empty vector (EV), RRAS or RRAS2 wild-type (WT), RRAS$^{Q87L}$ or RRAS2$^{Q72L}$. (**A**). Representative immunoblots from two independent experiments are shown. (**B**). Blots were quantitated by densitometry and the phospho/total protein ratios are shown. Results are the mean ± SEM. Statistical differences between groups was assessed by one-way ANOVA followed by Dunnett's multiple comparisons test. p<0.05 was considered statistically significant. Only p-values for significant comparisons are shown.

### Supplementary Figure 6

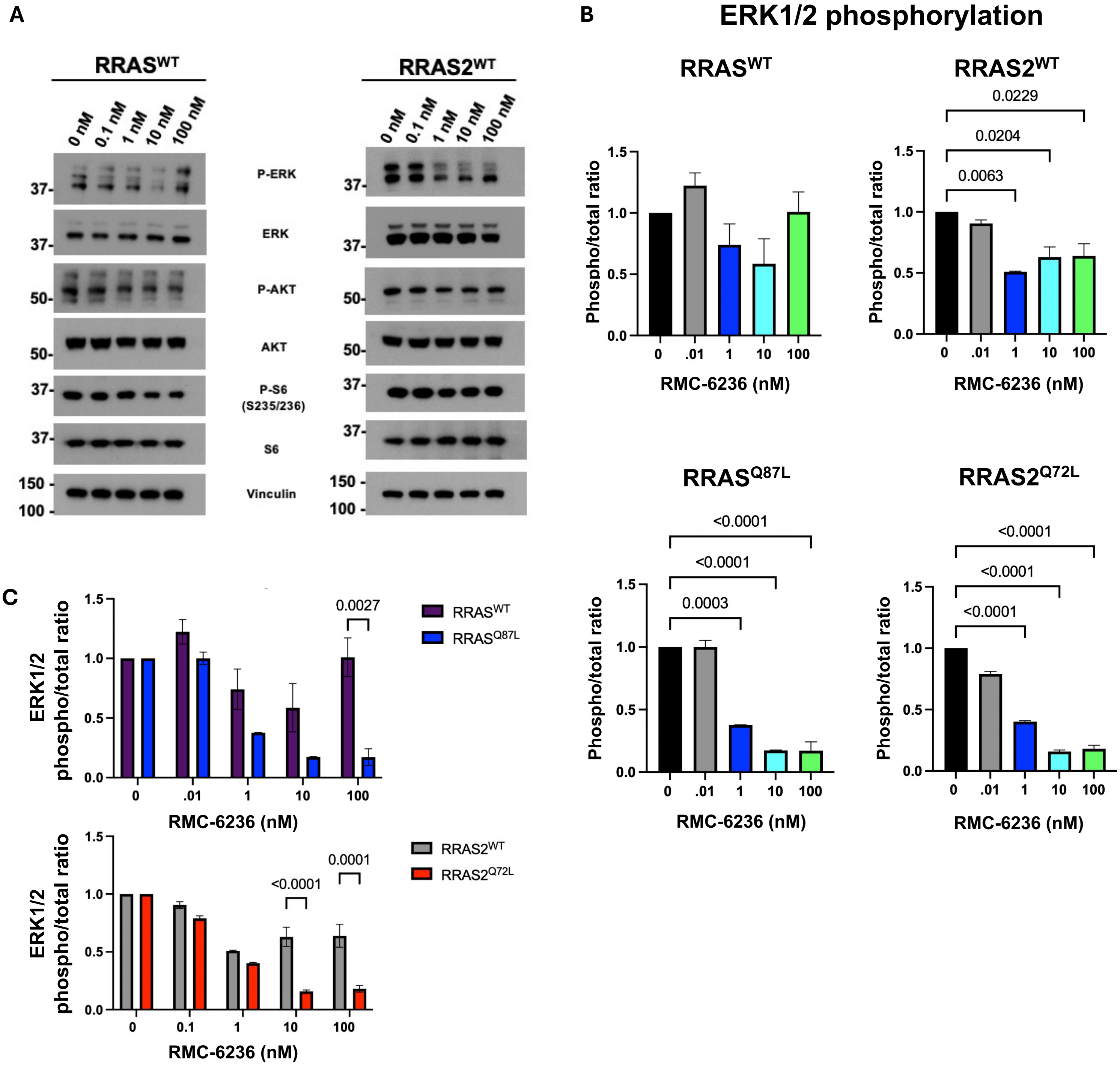


**Supplementary Figure 6.** Western blot analysis of extracts prepared from Ba/F3 cells expressing a wild-type RRAS or RRAS2 and treated with RMC-6236 for 90 min. (**A**). Representative immunoblots from two independent experiments are shown. (**B**). Blots for ERK1/2 phosphorylation quantitated by densitometry and the phospho/total protein ratios are shown. Comparisons made with one-way ANOVA followed by Dunnett's multiple comparisons test. (**C**) Comparison of relative ERK1/2 phosphorylation between RRAS/RRAS2 wildtype and mutants made with two-way ANOVA with Sidak's multiple comparisons test. Results represent the mean ± SEM. $p<0.05$ was considered statistically significant. Only p-values for significant comparisons are shown.

### Supplementary Figure 7

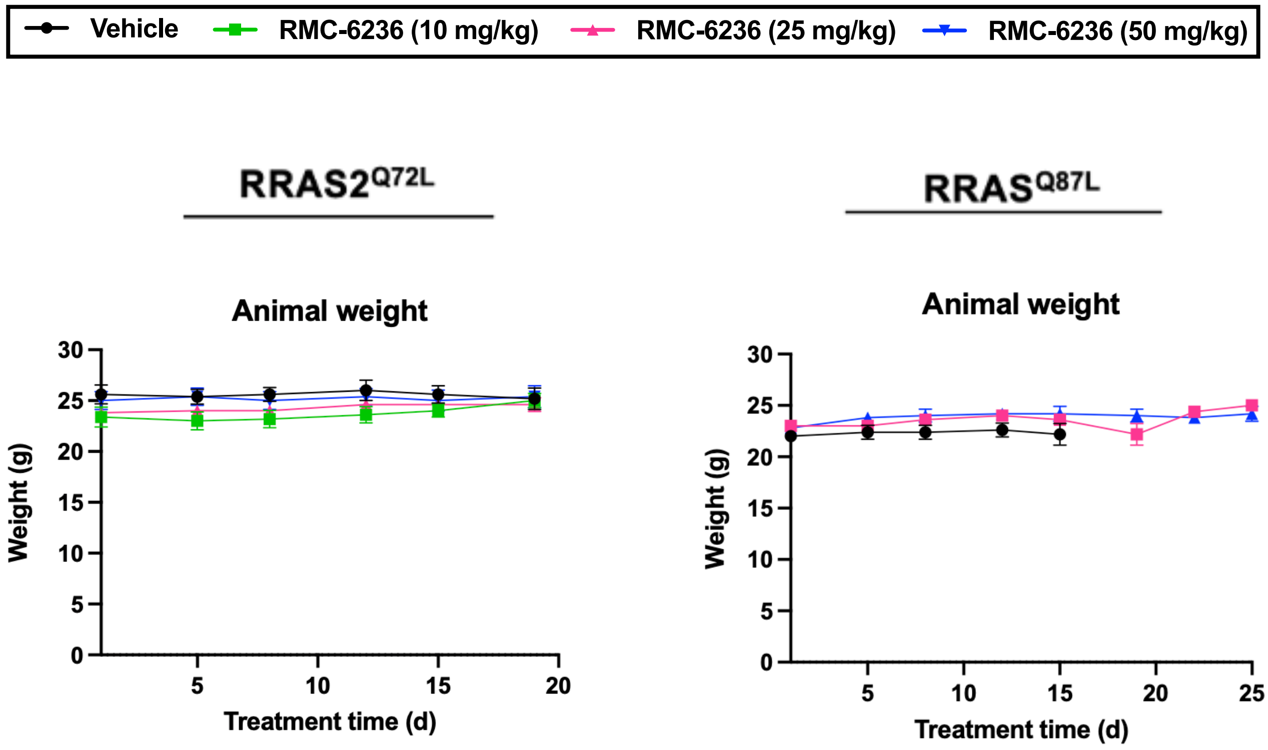


**Supplementary Figure 7.** Animal weight of immunocompromised NSG mice treated with RMC-6236 (vehicle, 25 mg/kg, or 50 mg/kg once daily for RRAS$^{Q87L}$-tumor bearing mice and vehicle, 10 mg/kg, 25 mg/kg, or 50 mg/kg once daily for RRAS2$^{Q72L}$-tumor bearing mice). The vehicle-treated RRAS$^{Q87L}$- tumor bearing mice were sacrificed at day 15 upon reaching a pre-determined tumor volume. Animal weight was measured twice weekly. There was no significant change in animal weight in any group through out the study.
